# An Atomistic view of Short-chain Antimicrobial Biomimetic peptides in Action

**DOI:** 10.1101/323592

**Authors:** Jagannath Mondal, Pushpita Ghosh, Xiao Zhu

## Abstract

Amphiphilic *β*-peptides, which are rationally designed synthetic oligomers, are established biomimetic alternatives of natural antimicrobial peptides. The ability of these biomimetic peptides to form helical amphiphilic conformation using small number of residues provides a greater synthetic advantage over the naturally occurring antimicrobial peptides, which is reflected in more potent antimicrobial activity of *β*-peptides than its naturally occurring counterparts. Here we address whether the distinct molecular architecture of short-chain and rigid synthetic peptides compared to relatively long and flexible natural antimicrobial peptides translates to a distinct mechanistic action with membrane. By simulating the interaction of membrane with antimicrobial 10-residue *β*-peptides at diverse range of concentrations we reveal spontaneous insertion of *β*-peptides in the membrane interface at a low concentration and occurrence of partial water leakage in the membrane at a high concentration. Intriguingly, unlike prototypical natural antimicrobial peptides, the water molecules leaked inside the membrane by these biomimetic peptides do not span entire membrane, as supported by free energy analysis. As a major advancement, this work brings into lights the key distinction in the membrane-activity of short synthetic biomimetic oligomers relative to the natural long-chain antimicrobial peptides.

## Introduction

The development of bacterial resistance to conventional antibiotics is a major concern towards public health. Antimicrobial peptides, which provide a natural defence against a large range of pathogens, including bacteria and fungi, are emerging as a sustainable substitute of antibiotics. The peptide-based antimicrobial agents are less prone to bacterial resistance than the natural antimicrobial agents and has got well-deserved prominence.^1–4^ However, serious issues with the naturally occurring antimicrobial peptides are that these natural peptides are susceptible to degradation by cellular proteases and lack specificity for microbial targets over host cells. In this regard, synthetic oligomers and synthetic peptides which adopt helical conformations, are coming up as a viable alternative to overcome these limitations of natural antimicrobial peptides.^5–8^ The present work focuses on a promising class of potentially novel biomimetic anti-microbial peptides, referred as *β*-peptides and provides key atomistic insights of the underlying mechanisms of their membrane-disruption capability, in comparison to their natural counterparts.

*β*-peptides are synthetic oligomers of *β*-amino acids and rationally designed to mimic and improve the biological activities of their natural counter parts, namely *α*-peptides (Figure 1). The backbone of a *β*-amino acid contains an additional carbon atom compared to *α*-amino acid backbone, which provides an additional site for side chain introduction and makes the *β*-peptides resistant to degradation by natural proteases.^9^ In this regard, synthetic oligomers and random copolymers of *β* amino acids^10,11^ have recently been shown to have potent antibacterial^12–14^ and antifungal^15,16^ activity. The system of our current interest is a 10-residue-length *β*-peptide sequence *β*Y-(ACHC-ACHC-*β*K)_3_ where *β*Y, *β*K and ACHC refer to *β*-homotyrosine, *β*-homolysine and trans-2-amino cyclohexyl carboxylic acid (see Figure 1a). The cyclic constraint of ACHC residues strongly stabilizes this significantly short *β*-peptide sequence into a so-called 14-helical conformation and the sequence arrangement provides an amphiphilic nature where hydrophobic ACHC and hydrophilic *β*-homolysine are globally segregated (see Figure 1b). The ability to form stable 14-helical amphiphilic conformation by only 10 *β*-amino residues provides this *β*-peptide sequence a greater synthetic advantage over numerous naturally occurring antimicrobial peptides such as magainin, melittin or LL37, which are relatively much longer in sequence-length.

**Figure 1:**
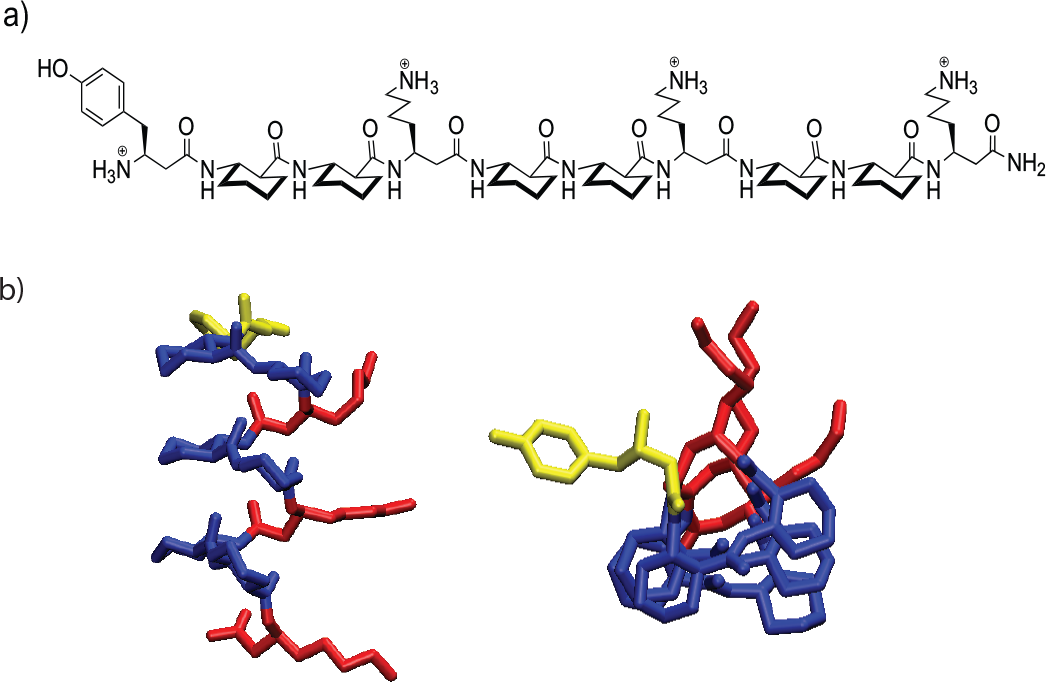
Structure and sequence of 10-residue antimicrobial *β*-peptide *β*Y-(ACHC-ACHC-*β*K)_3_ under current study. a) Chemical structure of *β*Y-(ACHC-ACHC-*β*K)_3_, b) Top view and side view of the three-dimensional structure of the same *β*-peptide in its stable 14-helical conformation.

Experimental investigations led by Gellman and coworkers^12^ have previously shown that this specific *β*-peptide sequence can impart strong anti-bacterial activity on both gram-positive and gram-negative bacteria at a very low minimum inhibitory concentration (MIC). The antimicrobial *β*-peptides were found to be comparable or more potent than the established antimicrobial natural peptides and were also found to be less hemolytic than the natural antimicrobial peptides. ^5^ The trends were similar for the anti-fungal activity of the current sequence of interest on *Candida albicans*^15^ with MIC value ranging between 16-32 *μ*g/ml. The *β*-peptide sequence of our interest (Figure 1) was also found to induce leakage of enzyme *β*-galactosidase from *Bacillus subtilis*,^13^ thereby suggesting membrane permeabilization by *β*-peptides. Nonetheless, while these experimental data provide convincing macroscopic evidence of antimicrobial activity of *β*Y-(ACHC-ACHC-*β*K)_3_, to the best of our knowledge, an atomistic insight towards the plausible mechanism of the antimicrobial activity of *β*-peptide, in comparison to natural antimicrobial peptides, has remained unexplored.

While multiple hypotheses exist^1–3^ in general to describe the principle steps leading to the membrane-disruption by antimicrobial peptides, central to all prevailing hypotheses is, the possibility of water pore formation across membrane by antimicrobial peptides.^17,18^ In fact many natural antimicrobial peptides namely magainin-H2, melittin are believed to form trans-membrane pore across the membrane bilayer.^19,20^ However, it is also believed that in general, there is no unique mechanism and it strongly depends on the specific antimicrobial peptides of interest.^21,22^ In these contexts, the pertinent question we ask in our current work: Does the synthetic peptide of our interest act by forming water-filling pore across the membrane? Or, does it disrupt the membrane integrity in so-called “carpet like”^3,4^ mechanism? The question is relevant mainly due to the significantly distinct molecular architecture of these short-chain rigid *β*-peptide sequence compared to its flexible and longer natural counterparts like magainin-H2 and melittin.

The present work, using a combination of multi-microsecond unbiased atomistic computer simulation and enhanced sampling of a 10-residue and 14-helical *β*-peptide sequence (presented in Figure 1) on membrane addresses these above-mentioned questions. By systematically varying the peptide/lipid (P/L) ratio in our simulations, we provide an atomistic account of the antimicrobial activity of the *β*-peptides on a negatively charged 1,2-dipalmitoyl-sn-glycero-3-phosphoglycerol (DPPG) lipid bilayer. The results so-obtained are further compared across a mixed bilayer of 1-palmitoyl-3-oleoyl-sn-glycero-2-phosphoethanolamine (POPE) and 1-palmitoyl-2-oleoyl-sn-glycero-3-phosphoglycerol (POPG) lipids which is known as a model bacterial-mimicking membrane. Using the force field specifically developed for *β*-peptides, ^23,24^ our simulation shows that at a lower P/L, the *β*-peptides glide through the aqueous media and bind spontaneously on the membrane head group region. At intermediate P/L, the *β*-peptides start to self-assemble and adsorption on the membrane-interface becomes aggregation-limited, with a very small subset of peptides ultimately adsorbing to the membrane-head group interface. At higher P/L, our long simulation trajectories reveal significant water leakage in the membrane interior, accompanied by membrane deformation. However, the water leakage induced by these antimicrobial *β*-peptides are not found to be trans-membrane-spanning. By subsequent enhanced sampling molecular dynamics simulations (see material and methods and Figure S1 for reaction coordinate) we confirm that the formation of a transmembrane pore would require these short peptides to align in a normal fashion with respect to the membrane interface and stretch to encompass the entire membrane, which is free energetically highly unfavorable. Nonetheless, apart from the observation of water leakage inside the membrane, at high P/L, these biomimetic peptides induce significant disorder in bilayer accompanied by membrane deformation due to strain imposed by the *β*-peptides. The work not only puts forward a working mechanism of the antimicrobial activity of the biomimetic peptides, but also elucidates key distinction in the mechanistic action between natural antimicrobial peptides and the short-chain synthetic oligomers.

## Results

### Membrane attachment of 5-peptides is aggregation-limited

Figure 2a) and 2b) respectively show the time profiles and the static density profiles of z-component of center of mass distance(C.O.M) of β-peptides and DPPG membrane center at different P/L ratios (see Material and Methods). As depicted in Figure 2, at low P/L ratio of 1:128 the *β*-peptide, initially present in aqueous media, gradually approaches the headgroup-water interface of the DPPG lipid bilayer and eventually, at a time scale of 500-800 ns, the *β*-peptide ultimately gets attached in the membrane interface of water and lipid head group as monomeric individuals. This is depicted by the density profile (see Figure 2b) of each individual *β*-peptide relative to that of the phosphate groups. Representative snapshots of the simulations, as shown in Figure 3, render a pictorial view of the membrane-partitioned state of the *β*-peptides at low to high P/L ratios. As the P/L ratio is increased, the *β*-peptides start to self-assemble and the approach of *β*-peptides towards the membrane interface becomes aggregation-limited. As illustrated by the time and density profiles and as rendered by the representative snapshots of the *β*-peptides in DPPG bilayer, at a medium P/L ratios equal to 4:128, our simulation result reveals that 2-3 copies of *β*-peptides self-aggregate in water and approach the membrane interface of DPPG bilayer as aggregate. The self-assembled structure of the *β*-peptides, with charged *β*-Lysine groups facing the negatively charged lipid bilayer prevent themselves in getting partitioned across the lipid head-group and water interface. This is evident from the density profiles (Figure 2b)) of *β*-peptides relative to the phosphate group of membrane at medium and higher P/L. We find that at very high P/L ratio, majority of the *β*-peptide copies can not get past that of the phosphate groups of DPPG membrane. A comparison with the representative snapshots in Figure 3 also confirms that at high P/L ratio, *β*-peptide copies, which remain as single individual entity, without getting self-assembled, can get past the phosphate group of DPPG membrane bilayer, while those which are part of an aggregate, remain limited to the membrane head group. A closer inspection of the snapshot of *β*-peptides in DPPG bilayer at higher P/L ratio further reveals that the *β*-peptide self-assembly is mainly driven by the hydrophobic association of ACHC *β*-amino acid residues and the aggregate remains stable due to favorable electrostatic interaction of the positively charged and exposed *β*-Lysine groups of the aggregate with the negatively charged head group of DPPG membrane. It is to be noted that the initial configuration of all simulation consists of all the *β*-peptides on same side of the membrane bilayer.

**Figure 2:**
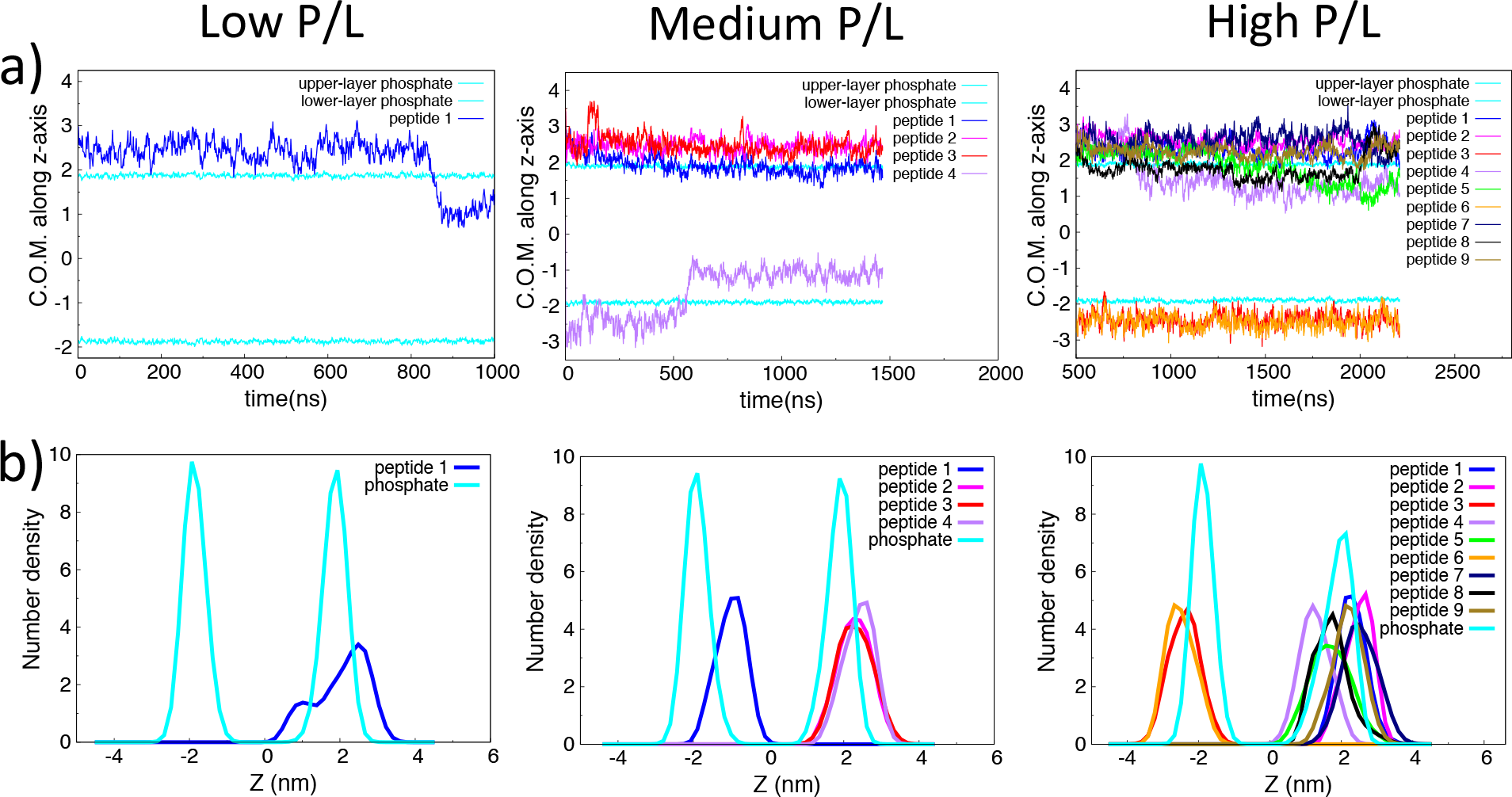
a) Time profile and b) density profile of Z direction of center of mass position of peptides relative to membrane center of mass of DPPG lipid bilayer at low, medium and high P/L ratio.

**Figure 3:**
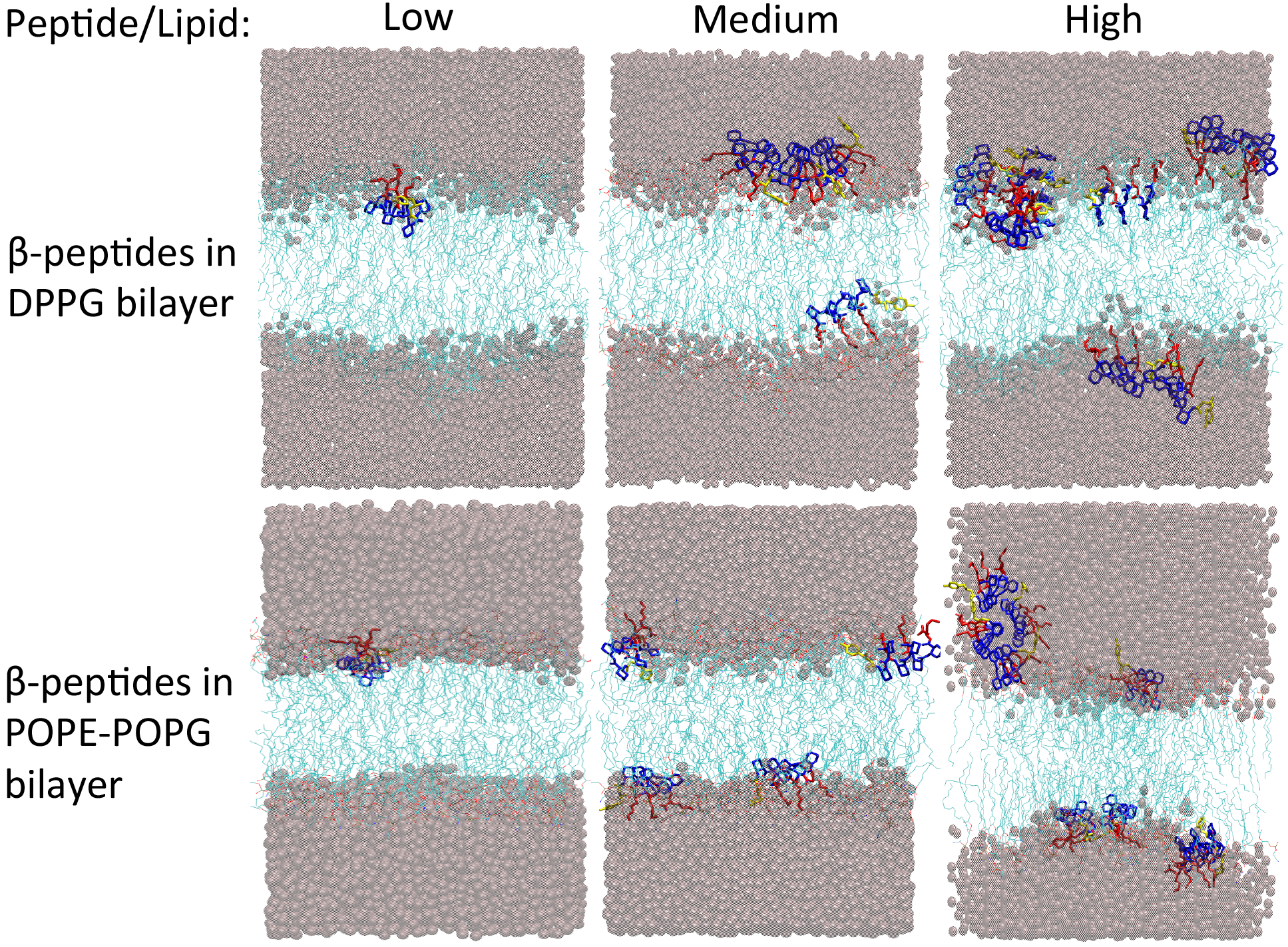
Representative snapshots of the *β*-peptide(s) and the interfacial water interacting with the membrane at a range of P/L ratio. Top: In DPPG bilayer and Bottom: In the mixed bilayer of POPE/POPG. Color code: lipid tails are cyan lines,head groups are red lines, water oxygen atoms are illustrated by pink spheres. The *β*-peptides are shown in licorice representation with hydrophobic ACHC residues in blue, *β*-lysine groups in red and *β*-tyrosines are in yellow. The initial configuration of all simulations have all copies of *β*-peptides on same side of the membrane.

As evident from the simulation snapshots, because of periodic boundary condition(PBC), the *β*-peptides are capable of arranging themselves on both the leaflets of the membrane in the equilibrated snapshot. However, we note that only a very small fraction of all the peptides, starting on the upper leaflet, gets adsorbed on the lower leaflet. Majority of the peptides stay on the upper leaflet and overall antimicrobial effects on the membrane is mainly guided by the peptides on the upper leaflet. As will be shown later, the membrane properties such as water density profile near lower leaflet, lipid tail order parameter of lower leaflet does not change significantly across P/L ratio, when compared with that near upper leaflet.

The self-aggregation propensity of specific amphiphilic *β*-peptide of our current interest is experimentally quite well known in aqueous media.^25–27^ The atomistic simulation of these *β*-peptides in presence of DPPG membrane interface presented in this article reveals that the self-assembly behavior of these *β*-peptides competes with their membrane-activity, making the membrane attachment process of the *β*-peptides aggregation-dependent. We note that our choice of DPPG membrane is mainly motivated by the fact that bacterial membrane is overall negatively charged and in fact negatively charged DPPG membrane has previously served as a suitable model membrane to explore antimicrobial property.^28^ However, it is known that PG-type lipid is only composed of 15 % of the entire phospholipid in *Escherichia coli* membrane and mostly the inner membrane.^29^ To further investigate the specific role of membrane-bilayer composition on membrane-activity and self-aggregation behavior of *β*-peptides, we carried out same set of simulations using a mixed membrane bilayer of POPE and POPG at a ratio of 5.3:1, which is usually considered as an optimum model of bacterial membrane.^30^ The density profiles of *β*-peptides in POPE/POPG mixed bilayer in Figure S2 in SI, show the same trend as in DPPG bilayer at low P/L ratio, with single copy of *β*-peptide reaching past the phosphate groups of the membrane. Corresponding snapshot at Figure 3 at low P/L ratio echoes the same trend in both DPPG and POPE/POPG bilayer, with the *β*-peptide getting partitioned across the bilayer. On the other hand, at medium P/L ratio, representative snapshots of the *β*-peptides in POPE/POPG bilayer show that the *β*-peptides actually approach the bilayer as single monomeric individual and spontaneously get partitioned across the bilayer interface past the phosphate groups. The absence of self-assembly of the *β*-peptides in presence of POPE/POPG bilayer (at 5.3:1 ratio) illustrates a contrasting picture of *β*-peptides in DPPG bilayer at intermediate P/L ratio. We believe that significantly reduced negative charge density of lipid head groups in POPE/POPG bilayer compared to DPPG bilayer destabilizes any preformed *β*-peptide aggregate and results in the dispersal of the aggregates on the membrane interface. However, as seen in the representative snapshots in Figure 3, higher P/L ratio promotes stronger *β*-peptide self-assembly and hence the *β*-peptides adsorb on the membrane as aggregate in both DPPG membrane and POPE/POPG mixed membrane. In other words, at a very high P/L ratio, *β*-peptides respond almost similarly to both DPPG and POPE/POPG bilayers. Hence, as a common point of interest, at high P/L ratio, we hereafter focus on the action of *β*-peptides towards single DPPG membrane only. The observed self-aggregation propensity of *β*-peptides, specially in presence of negatively charged membranes in the current work, suggests that *β*-peptides might have the potential of triggering bacterial agglutination as part of its mechanism of action as previously noted in some antimicrobial peptides and amyloids.^31,32^ However, this certainly needs to be tested by new experiments in future.

### *β*-peptides remain parallel and helical upon membrane adsorption

The angular distribution of the *β*-peptides relative to the surface-normal of membrane sheds light on the fashion of membrane-attachment of the *β*-peptides. This is illustrated in Figure S3 (see SI). Our simulation results show that at very low P/L ratio, the peptide individually attaches to the membrane interface (Figure 2b)), the angular distributions with the z-axis tend to peak around 80-90 degree implying the membrane-attachment of these *β*-peptides in a parallel fashion (Figure S3 in SI). As the concentration of *β*-peptide increases, the membrane attachments become aggregation limited which also influences the relative orientation of the *β*-peptides with respect to the membrane interface. As depicted in Figure S3, in case of medium to high P/L ratios, those *β*-peptides which could get attached to the membrane interface individually without getting self-aggregated in aqueous media, maintain a parallel orientation with the membrane interface. On the other hand, the *β*-peptide copies which get self-aggregated in aqueous media and can not partition themselves across the membrane interface, retain an isotropic distribution typical of aqueous media. The spontaneous propensity of *β*-peptides attaining a parallel orientation with the membrane interface after attachment can be rationalised by their amphiphilic nature. The *β*-peptides attach to the membrane-surface in a parallel fashion so that the hydrophobic ACHC groups can remain in touch with the hydrophobic lipid tails and charged *β*-Lysine residues can optimally interact with the membrane polar head groups.

*β*-peptides are known for their high stability of 14-helical conformation in aqueous media. ^33^ It is therefore a key question how the interaction with the membrane interface can influence the secondary structure of the *β*-peptides. Figure S4 in SI compares the distribution of root mean squared deviation of C-alpha atoms from the ideal helical structure of these *β*-peptides after membrane-attachments. We find that the secondary structure remains 14-helical even after membrane insertion. This is consistent with the previous experiments by Gellman and coworkers^12,15^ suggesting that the *β*-peptides maintain same signature of secondary structure both in membrane and in aqueous media. The observed retention of the helical nature by these short-chain *β*-peptides both in aqueous solution and in membrane interface are quite distinct from some of the well-studied natural antimicrobial peptides. Melittin and magainin-H2 ^19,20^ for example do not require strong helical architecture yet demonstrate significant antimicrobial activity. As we will find in the next sections, the propensity of retaining the 14-helical structure by these short-chain helical *β*-peptides in the membrane environment, compared to the flexibility displayed by natural antimicrobial peptides, have significant implication in the membrane-behavior of these synthetic oligomers.

### *β*-peptides at a high P/L ratio induce partial water leakage in membrane

The most interesting observation of our current work is that at high P/L ratio, water molecules partially leak through the upper leaflet of the membrane. As presented by the water density profile in Figure 4a) near upper leaflet, we find that while there is negligible water density inside the membrane at low P/L ratio, the number of water molecules gradually increases with higher peptide concentrations, and with a considerable number of pore waters within the membrane center at a very high P/L ratio of 9:128. At this high peptide to lipid ratio of 9:128, the water number density is significantly higher in the membrane interior compared to that at a lower P/L ratio. The representative snapshots shown in Figure 4b) and 4c) also render a pictorial view of the water leakage inside the membrane interior. Interestingly, we find that water molecules translocating inside the membrane interior closely follow the *β*-peptides which had got adsorbed in the membrane. The observation implies that these *β*-peptides, while getting partitioned in the membrane-water interface, also bring water molecules along with them, thereby leading to water leakage. However, intriguingly, we find that these water leakage induced by the *β*-peptides are not membrane-spanning in nature. We reiterate that a P/L ratio of 9/128 might seem relatively higher than what is generally being used for natural antimicrobial peptides such as magainin-H2 or mellittin. However, we here emphasize that these *β*-peptides are significantly shorter (only 10 residues) in chain-length and lower in molecular weight than that of magainin-H2 (23-residue), melittin (27-residues) or LL37 (37-residues) and hence for a similar P/L ratio, the concentration of the amphiphilic *β*-peptides will be much lower than that of typical antimicrobial peptides. Our specific observation of water pores not being transmembrane-spanning in the case of *β*-peptides is quite distinct compared to the two well-studied natural antimicrobial peptides, melittin and magainin-H2. Previously, the formation of a transmembrane-spanning toroidal pore has been well established for melittin.^20,34^ It has also been suggested that multiple copies of melittin exercise sufficient flexibility in lining along the water pores. Similar observation of toroidal pore, albeit disordered, has also been made for magainin-H2.^19^ Taken together, these comparisons allude to a different mechanism of action for the *β*-peptides compared to the natural peptides and calls for a physical interpretation of these suggested differences in the case of these synthetic oligomers.

**Figure 4:**
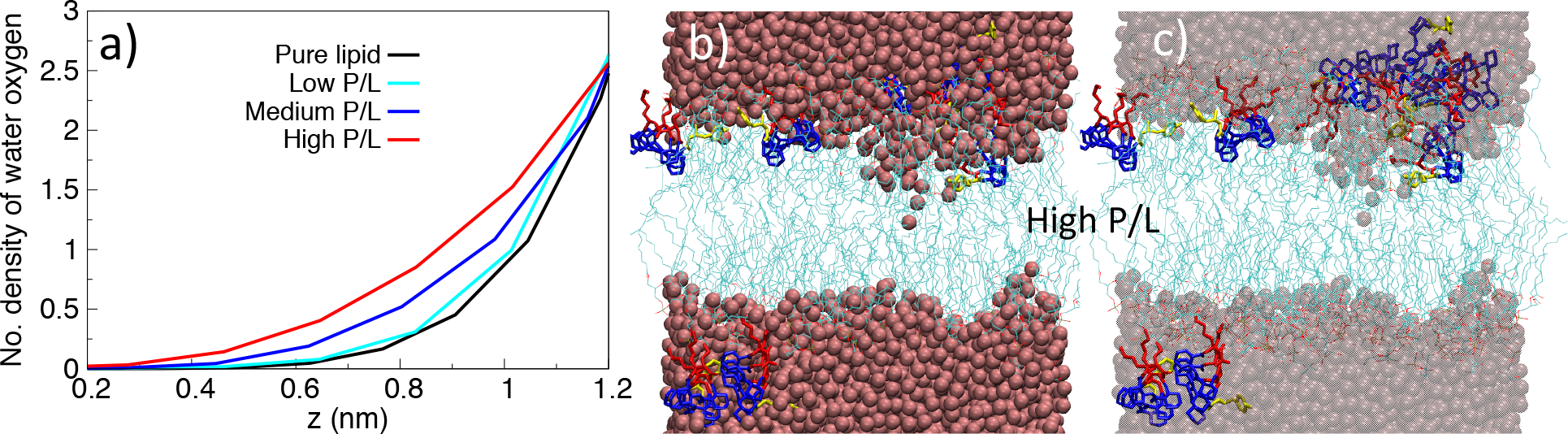
a): Comparison of the number density of the water molecules of DPPG bilayer as a function of P/L ratio. The results are shown only for upper leaflet of the bilayer. b) Representative snapshot of water leakage inside the membrane for P/L ratio of 9/128 in DPPG bilayer. Color code: lipid tails are cyan lines,head groups are red lines, water oxygen atoms are illustrated by pink spheres. The *β*-peptides are shown in licorice representation with hydrophobic ACHC residues in blue, *β*-lysine groups in red and *β*-tyrosines are in yellow. c) Same figure as in b) but the water molecules are blurred so that *β*-peptides are shown with clarity

### Formation of trans-membrane water channel by short-chain *β*-peptides is free energetically unfavorable

To investigate further why these peptide-induced water leakages are not membrane-spanning in nature, we perform umbrella sampling simulation, an enhanced sampling technique, to understand the underlying free-energetic origin. The details are provided in Material and Methods. Specifically, we choose a three-member stable cluster of the *β*-peptides that is partitioned at the membrane-water interface and the umbrella sampling is performed using the z-component of the distance of separation of the center of mass of the cluster and the membrane as the designated reaction coordinate (*ξ*) (see Figure S1 in SI for a schematic representation of the reaction coordinate). The resulting free energy profile as a function of collective variable, as depicted in Figure 5, provides key insights on the interaction of the membrane with the peptide and on the plausibility of transmembrane water pore formation. The global minima of the free energy profile corresponds to the aforementioned equilibrium structure of subset of peptides being adsorbed at the interface at *ξ*=1.5 nm, with water being partially translocated in the membrane. However, as the value of of ξ decreases, the peptide cluster gradually approaches towards the center of the membrane and we find that the free energy monotonically increases. The maxima of the free energy profile at *ξ*=0, in fact corresponds to a conformation in which the peptides are vertically stretched out across the membrane bilayer accompanied by the formation of the transmembrane pore with water being lined up encompassing the entire bilayer. This suggests that the formation of so-called transmembrane pore is free energetically highly unfavorable in the case of these short-chain *β*-peptide oligomers. Fig. S5 in SI text provides a quantitative illustration of this observation, by projecting the free energetics along both *ξ* and Number of waters present within the membrane(*N*_*w*_), as computed from trajectories of umbrella-sampling windows. We find that while the free energy minima near the interface at *ξ*=1.5 nm is associated with considerable number of waters, the peptide positioned in the membrane interior (corresponding to smaller ξ) might be associated with very large number of water but at the expense of very high free energy. A multi-dimensional projection of the free energy profile along *ξ* direction and the angle of orientation with the z-axis, as shown in top right of Figure 5, provides further insights on the prohibitiveness of transmembrane pore formation by these antimicrobial *β*-peptides. We find that the *β*-peptides are required to have a vertical orientation with the membrane surface (angle with the z-axis being between 0-30 degree) to induce a membrane-spanning pore at smaller *ξ*, which is free energetically highly unfavorable. This can be attributed to the fact that the vertical orientation with the membrane surface exposes the hydrophilic βLysine residues to the hydrophobic lipid tails which is free energetically very expensive. Moreover, the inherently rigid structure and short chain length makes it thermodynamically unfavorable for the *β*-peptides to stretch across the membrane, rendering the transmembrane orientation of the accompanying water highly unlikely. Previous free energy estimations for natural antimicrobial peptide melittin^35^ have suggested a free energy barrier between surface-bound and transmembrane state of melittin and it has been found that the presence of multiple copies of melittin can stabilize the transmembrane structure. However, this is majorly due to the flexible helicity of melittin which provides key impetus for a transmembrane orientation. Together, these results provide a clear rationale for this distinct behavior of the antimicrobial *β*-peptides in membrane environment and points towards the short and rigid molecular architecture of the *β*-peptides as a key factor in deciding its nature of membrane action.

**Figure 5:**
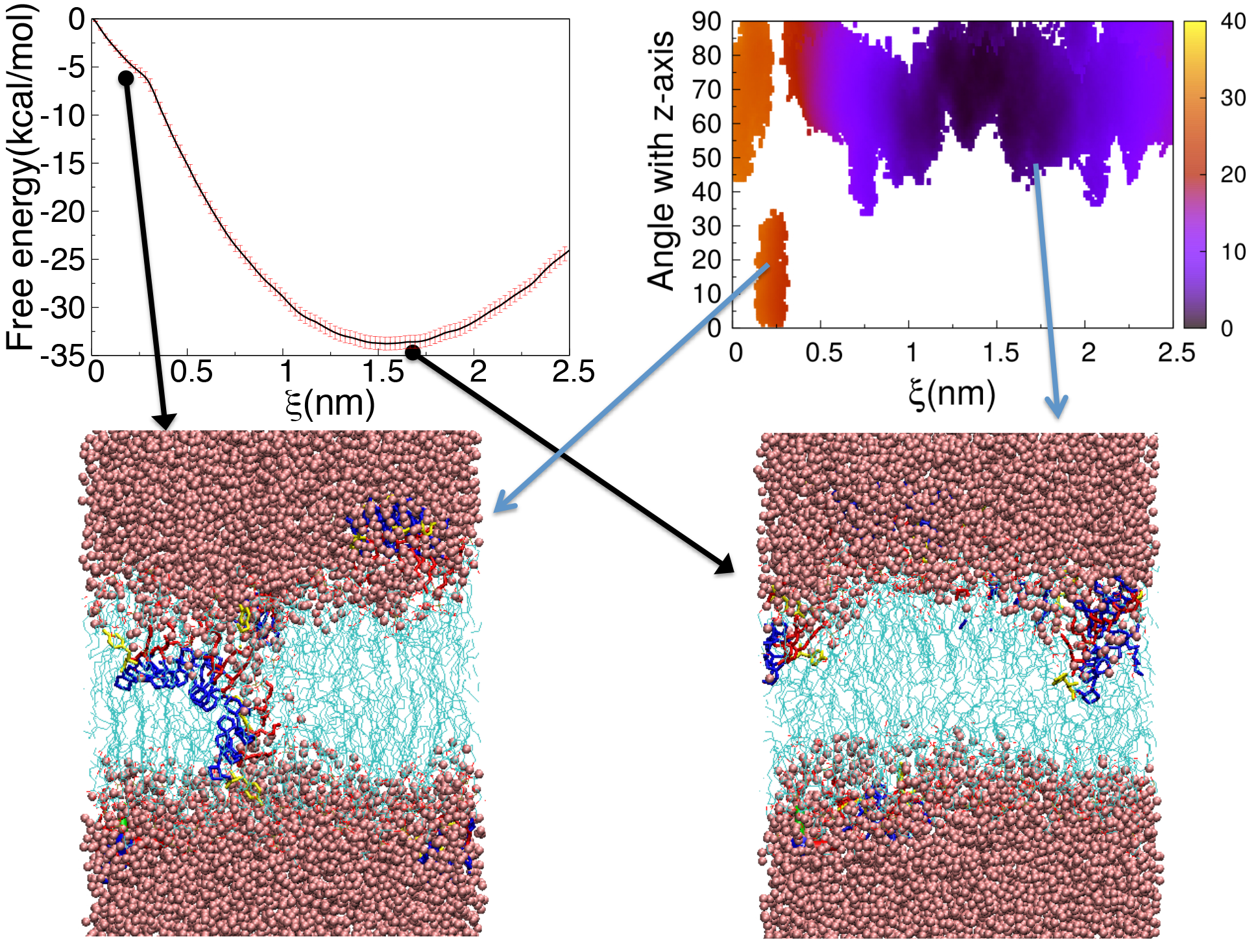
Top Left: Free energy profile of internalization of a three-peptide aggregate with DPPG bilayer as a function of z component of peptide-membrane separation. Top left: Two-dimensional Free energy profile of the association of a three-peptide aggregate with the DPPG bilayer as a function of z-component distance of peptide-membrane separation and angle of the peptide-cluster orientation relative to the membrane normal. Bottom: Representative snapshots near minima and maxima of free energy profile. Color code: lipid tails are cyan lines, head groups are red lines, water oxygen atoms are illustrated by pink spheres. The *β*-peptides are shown in licorice representation with hydrophobic ACHC residues in blue, *β*-lysine groups in red and *β*-tyrosines are in yellow.

We note that the choice of a specific cluster for umbrella sampling is majorly guided by the observation of a cluster, which remains stable over a significant period of simulation time. While we have chosen a single cluster to perform the free energy calculation of peptide internalization, we note that the purpose of this umbrella sampling simulation has to be considered as a test of hypothesis of possibility of pore formation by the beta-peptide aggregates. The free energy profile computed from umbrella sampling paints a qualitatively consistent picture with the observation from unbiased multi-microsecond simulations: while the beta-peptides can induce partial water leakage inside the membrane, the chances of formation of trans-membrane pore are remote. It is quite possible that the free energy profile reported in the work can have quantitative variation with respect to choice of cluster. However, we believe that the qualitative picture obtained in the current study will remain unchanged.

### *β*-peptides at a higher concentration disrupt membrane integrity

The antimicrobial agents are known to influence the mechanical integrity of the membrane bilayer. To investigate the effect of these *β*-peptides on the membrane, we compute the deuterium order parameter of the lipid tails and the two-dimensional density profile of the lipid molecule on the membrane-attaching leaflets of the bilayer. As depicted in Figure 6(a), the value of the order parameter increases with the increase of peptide concentration suggesting that peptide attachment on the top of the bilayer retards the motion of the lipid molecules beneath it. This trend is quite similar to that reported by Vemparala and coworkers,^36^ albeit for a different antimicrobial peptide. As a consequence of antimicrobial activity of these *β*-peptides, inhomogeneity in the lateral density profile of the membranes is also observed as the peptide/lipid ratio is gradually increased. As is evident from the density profiles in Figure 6(b), with increasing peptide concentration, the self-aggregation of the *β*-peptides causes significant fluctuation of density in the membrane morphology. The stress induced by the self-assembled *β*-peptides induces deep trough in the membrane with certain regions of the membrane having larger height. The surface area per lipid also reduces considerably with the increase of peptide concentration (See Table S1 in SI). The overall trend in bilayer thickness and membrane surface area points towards strong antimicrobial action of these short-chain *β*-peptides. Finally, we note very weak variation in the water density profiles and lipid-tail order parameters of the lower leaflets with increasing peptide concentration(Figure S6 in SI). These results collectively suggest that the perturbation in the lower leaflet of the lipid bilayer is minimal and the overall antimicrobial effect of *β*-peptides is majorly induced by the *β*-peptide adsorption in the upper leaflet only.

**Figure 6:**
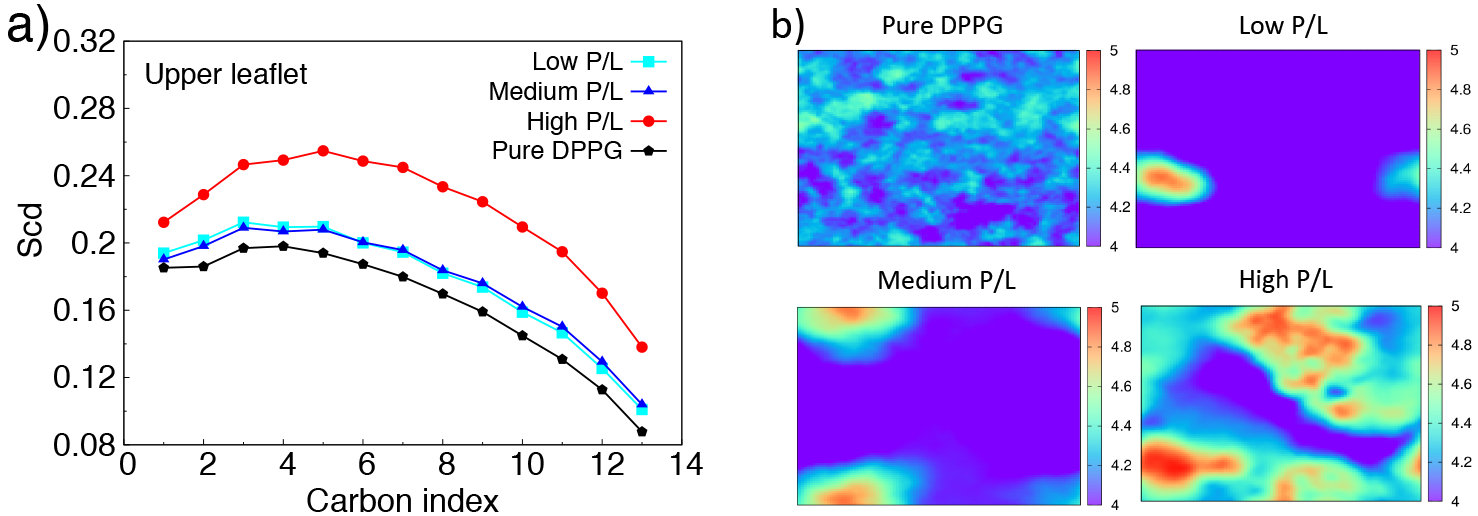
a) Comparison of Deuterium order parameter(Scd) of the DPPG upper leaflet at different P/L ratio. b) Comparison of bilayer thickness profile of the DPPG leaflet at different P/L ratio.

## Discussion

Unravelling the mechanistic action of antimicrobial agents at an atomistic resolution has always been desirable from both experimental and computational perspectives. Specifically, it remains more intriguing to decipher the membrane-disrupting mechanism for amphiphilic *β*-peptides which manifest strong antimicrobial properties despite being much shorter in chain-length and lower molecular weight than their natural counterparts. Towards this end, our current microsecond computer simulation at an atomistic length scale closes an important gap towards understanding the antimicrobial activity of these short-chain synthetic oligomers. Unlike precedent computer simulation studies, which have explored mainly membrane-attachment of peptides at a low Peptide/Lipid ratio or which have started with manually preformed specific transmembrane pore,^21,22^ our current work takes a step forward and investigates the thermodynamic feasibility of membrane disruption of these rigid 10-residue-long *β*-peptide foldamers with extensively varied P/L ratios. The present work brings out key differences between the rod-like short-chain synthetic *β*-peptides and the long-sequenced flexible natural peptides in their interaction with the membrane. As a key result, we observe spontaneous water leakage in the membrane at a high P/L ratio. Our work, using enhanced computer simulation approach, also provides a quantitative free energetic account of the prohibitiveness of transmembrane pore formation by these short-chain *β*-peptides as opposed to the observed partial water leakage. The ability to simulate the spontaneous formation of water pore inside the membrane interior via multi microsecond atomistic computer simulations also speaks highly in favor of specially designed CHARMM-compatible parameterized forcefield of *β*-peptides.^23,24^ Our work puts forward a distinctive case regarding the antimicrobial activity of non-natural oligomers and compares it with established natural peptides. Overall, our work unequivocally shows that, unlike natural peptides, which impart their antimicrobial activity via formation of trans-membrane water-pore across the membranes, these short and rigid antimicrobial *β*-peptides have a distinct mechanistic action: A joint effort by induction of partial water-leakage in the membrane coupled with aggregation-driven membrane deformation by these *β*-peptides is required to disrupt the membrane integrity.

The current work opens up multitudes of opportunities towards understanding of the mechanistic aspects of the membrane-disruption capability by both natural and biomimetic macromolecules. The large scale membrane deformation via curvature formation by many charged peptides^37^ including *β*-peptides ^38^ is of utmost interest and in this regard the development of a coarse-grained model for *β*-peptides is currently underway. Moreover, recent works by Gellman and coworkers on exploring random copolymeric *β*-peptides as cost-effective and efficient antimicrobial agents are being seen as major paradigm shifts.^39–41^ These works suggest a departure from the preconceived notion of helical propensity as a key pre-requisite for the manifestation of antimicrobial activity and open up possibilities of venturing into atomistic view of these random copolymeric antimicrobial agents. The significant aggregation propensity of *β*-peptides in presence of membrane as observed in the current work also partly insinuates that *β*-peptides might have the potential of triggering bacterial agglutination as part of its mechanism of action as found in some antimicrobial peptides and amyloids, ^31,32^ which certainly warrants future experiments to test the idea.

Finally, pore formation capability by antimicrobial peptides is currently being looked at as only one of the multiple facets of the overall realm of action of these membrane-disrupting agents.^31,42^ The membrane active natural peptides, such as melittin, have been found to induce lipid extraction from membranes,^43^ a critical process for many biological events. In this regard, specific segment of peptides, cholesterol and methylation levels of phospholipids among many others have been found to play an important role in membrane fragmentation.^44,45^ These works unravel new prospects of future investigations.

## Materials and Method

Due to the complex nature and composition of bacterial membrane, in this work we focus on a relatively simpler lipid bilayer model which is used as a good membrane-mimic for in vitro study. Majority of our simulations has been performed by the membrane constituted by a negatively charged bilayer using a total of 128 DPPG lipid molecules. The use of DPPG lipid bilayer is mainly due to the fact that bacterial membrane are overall negatively charged and DPPG membrane has previously been used as a model membrane to explore antimicrobial property. ^28^ The utilization of DPPG bilayer also provides us a good reference for exploring the membrane activity of the 5-peptides on a purely negatively charged bilayer. Nonetheless, we have also compared significant part of our results using a mixed bilayer of POPE and POPG at a ratio of 5.3:1, which is currently being considered as an optimum bacterial membrane model.^30^ The lipid bilayers were first assembled along with water molecules (around 7200 molecules) using CHARMM-GUI web server^46^ and charge neutralized using K^+^ ions. The charge neutralized lipid bilayer was then energy-minimized and equilibrated for 200 ns (simulation schemes are detailed later). Three different systems were set up by independently inserting 1, 4, and 9 copies of *β*-peptides respectively in the aqueous media of the equilibrated DPPG bilayer and subsequently removing the overlapping water molecules. We will refer these systems with the peptide to lipid ratio of 1/128, 4/128 and 9/128 as “Low", “Medium" and “High" P/L. It is worth mentioning that in all our simulations, all *β*-peptide copies were initially placed on same side of the membrane in the aqueous media. However, the periodic boundary condition might allow the peptides to remain adsorbed on both leaflets. But, as we will find from our simulation results, the net density of peptides on upper leaflet was be found to be always higher than that at lower P/L ratio and the membrane properties near the lower leaflets were largely unchanged across P/L ratio. We also note that similar protocols of periodic boundary conditions in all three directions have been used in many previous works.^19,20^ One of the rationales behind using periodic boundary condition in these works has been that peptides can be found to remain adsorbed asymmetrically or symmetrically on both leaflets of bilayers.^34^

To account for the residual positive charges introduced by the addition of the *β*-peptides (each having three *β*-lysine groups) in the aqueous media of the membrane, appropriate number of K^+^ ions were removed so that each of the final systems are rendered charge-neutral. Figure S7 represents the initial configuration of the simulation in two cases: namely, single copy of *β*-peptide in the aqueous phase of the membrane (Figure S7 a)) and nine copies of *β*-peptides in the aqueous phase of the membrane, all in the same side of the bilayer (Figure S7 b)). We note that a P/L ratio of 9/128 of *β*-peptide to lipid molecules might apparently seem to be much higher than that usually employed in computer simulation for natural antimicrobial peptides such as magainin, melittin or LL37. However, we need to take into account the fact that the specific *β*-peptide of our interest is only 10-residue length oligomer in contrast to relatively longer sequence-length and higher molecular weight of typical natural antimicrobial peptides (23 residues for magainin and 27 residue for melittin and 37 residues for LL37). Hence, the overall concentration of the *β*-peptides used in the current work is at par with their relatively longer natural counterparts.

CHARMM36 forcefield parameters^47^ were employed to model lipid molecules, as these are known to provide a tensionless bilayer at ambient temperature and pressure. The CHARMM forcefields, specifically parameterized for these *β*-peptides by Zhu et al.,^23,24^ and previously used by Mondal et al^48^ was employed here to model the *β*-peptides. Finally, the water molecules are modeled by CHARMM-TIP3P water model^49^ due to its compatibility with the parameters of lipid molecules. CHARMM^47^ parameters were also used for the parameters of ions. Each of these systems with *β*-peptide(s) and lipid molecules were simulated until majority of the peptide copies get attached to the membrane surface. The simulation length, in this process, was ranged between 1 to 3 microsecond for each trajectory. All simulations were performed using a time step of 2 femtosecond and each simulation was repeated multiple occasions by varying the velocity seeds.

As will be discussed in the results later, our simulated trajectory with P/L of 9/128 reveals spontaneous water leakage inside the membrane i.e. water molecules spontaneously leak through the membrane interior. To investigate the free energetics of water pore formation in the system composed of 9 *β*-peptides and DPPG lipid bilayer, apart from equilibrium simulations, we performed additional umbrella sampling simulations. Towards this end, we used the z-component of the distance between the center of mass of the membrane and the center of mass (C.O.M.) of a cluster of three copies of *β*-peptides that were located at the membrane-water interface, as a reaction coordinate designated as *ξ* (For the illustration, see Figure S1 in Supporting information(SI)). The choice of this reaction coordinate is mainly motivated by the previous observation^50^ that pulling a phosphate group to the membrane center triggers the formation of a water pore, primarily because the charged phosphate group drags water inside the membrane. Hence, formation of water pore has been always considered a result of peptide insertion inside membrane. Similar reaction coordinate has been used in past^50–52^ with success and has provided very useful insights. We have employed umbrella sampling scheme for 26 windows corresponding to the values of *ξ* ranging from 0 to 2.5 nm, at an interval of 0.1 nm. The initial configuration corresponding to *ξ* = 0.3 to 2.5 nm were partly obtained from equilibrium simulation trajectory involving P/L of 9:128 and were partly obtained from steered molecular dynamics simulation. Each of the windows were subjected to a harmonic restraint of force constant 2000 kJ/mol/nm^2^ and then sampled for 150 ns (hence a total sampling of 3.9 microsecond). Finally, Weighted Histogram Analysis Method (WHAM)^53^ has been implemented to obtain underlying potential of mean force from the umbrella sampled time-series data. Bootstrapping error analysis has been performed to estimate the statistical uncertainties associated with the free energy profile. As will be discussed in the results section, we have also projected the free energy landscape along other dimensions, namely number of water in the membrane-interior (*N*_*w*_) and angle of peptides with membrane-normal by reweighing the joint probability distribution with the one-dimensional umbrella-sampled free energy along *ξ*. However, a more quantitative probe of pore formation might require explicit incorporation of membrane-water density as accompanying reaction coordinate in the umbrella sampling approach, which will be considered in future.^54^

All atomistic simulations were performed using Gromacs-5.0.6 software package.^55^ Each of the simulations was first subjected to an energy-minimization and subsequently, classical Molecular dynamics was performed at constant pressure of 1 bar and constant temperature of 323 K. Each of the components (*β*-peptides, lipids and water molecules) were also coupled separately with the thermostat. The temperature was maintained at the desired value by employing Nose Hoover temperature coupling scheme^56,57^ using a coupling constant of 1 ps. A semi-isotropic pressure coupling using Parrinello-Rahman protocol^58^ was implemented to maintain the desired pressure of 1 bar. The pressure coupling constants and compressibilities along xy and z directions were 5 ps and 4.5 × 10^5^ bar^−1^ respectively. In all simulations, center of mass motions were removed every 100 steps for *β*-peptides, lipids and water molecules individually. Verlet cutoff^59^ schemes were implemented for Lennard-Jones interaction extending upto 1.2 nm while particle mesh Ewald schemes were implemented for treating electrostatic interactions. All bond lengths involving hydrogen atoms of the protein and the lipids were constrained using the LINCS^60^ algorithm and water hydrogen bonds were fixed using the SETTLE approach.^61^ Simulations were performed using the leapfrog integrator with a time step of 2 fs and initiated by randomly assigning the velocities of all particles from a Maxwell-Boltzman distribution.

To quantify the order of lipid bilayers, second rank order parameter^62^ *P*_2_ = 1/2(3*cos*^2^(*θ*) − 1) was computed for consecutive bonds, with *θ* being the angle between the direction of the bond and the normal to the bilayer. Perfect alignment with the bilayer normal is indicated by *P*_2_ = 1, perfect anti alignment with *P*_2_ = −0.5, and a random orientation with *P*_2_ = 0. The lateral density profile and the area per lipid of the membranes (by considering the presence of the *β*-peptides) were also computed using “GRIDMAT MD” program.^63^

## Acknowledgments

This work was supported by computing resources obtained from shared facility of TIFR Center for Interdisciplinary Sciences, India. JM would like to acknowledge research intramural research grants obtained from TIFR, India, Ramanujan Fellowship and Early Career Research funds provided by the Department of Science and Technology (DST) of India (ECR/2016/000672). Part of the work was carried out in San Diego supercomputing resources provided by XSEDE (TG-CHE160057) to JM and XZ. PG would like to acknowledge the financial support by INSPIRE Faculty Grant (DST/INSPIRE/04/2015/002495) from DST of India. JM thanks Dr. Heejun Choi for a critical reading of the manuscripts and useful comments.

## Supporting Information (SI)

Graphs illustrating the reaction coordinate chosen for free energy simulation, *β*-Peptide density profiles in POPE/POPG bilayer, angular distribution of peptides with membrane surface and root mean squared deviation of *β*-peptides from the helical structures, projection of free energy profiles along distance and membrane-water number, bilayer properties in lower-leaflets, table containing Area-per-lipids of DPPG bilayer at different peptide to lipid ratio, Starting configuration of simulation (PDF).

## Supporting Information

**Table S1.** Area-per-lipids of DPPG bilayer at different peptide to lipid ratio

| Upper leaflet |  |
| --- | --- |
| System | Area-per-lipid ($\text{\AA}^2$) |
| Pure DPPG | $65.31 \pm 1.00$ |
| P/L=1/128 | $61.13 \pm 1.84$ |
| P/L=4/128 | $55.22 \pm 1.84$ |
| P/L=9/128 | $40.97 \pm 1.14$ |
| Lower leaflet |  |
| System | Area-per-lipid( $\text{\AA}^2$) |
| Pure DPPG | $65.32 \pm 0.90$ |
| P/L=1/128 | $65.95 \pm 1.54$ |
| P/L=4/128 | $61.11 \pm 1.79$ |
| P/L=9/128 | $58.29 \pm 1.21$ |

**Figure S1:**
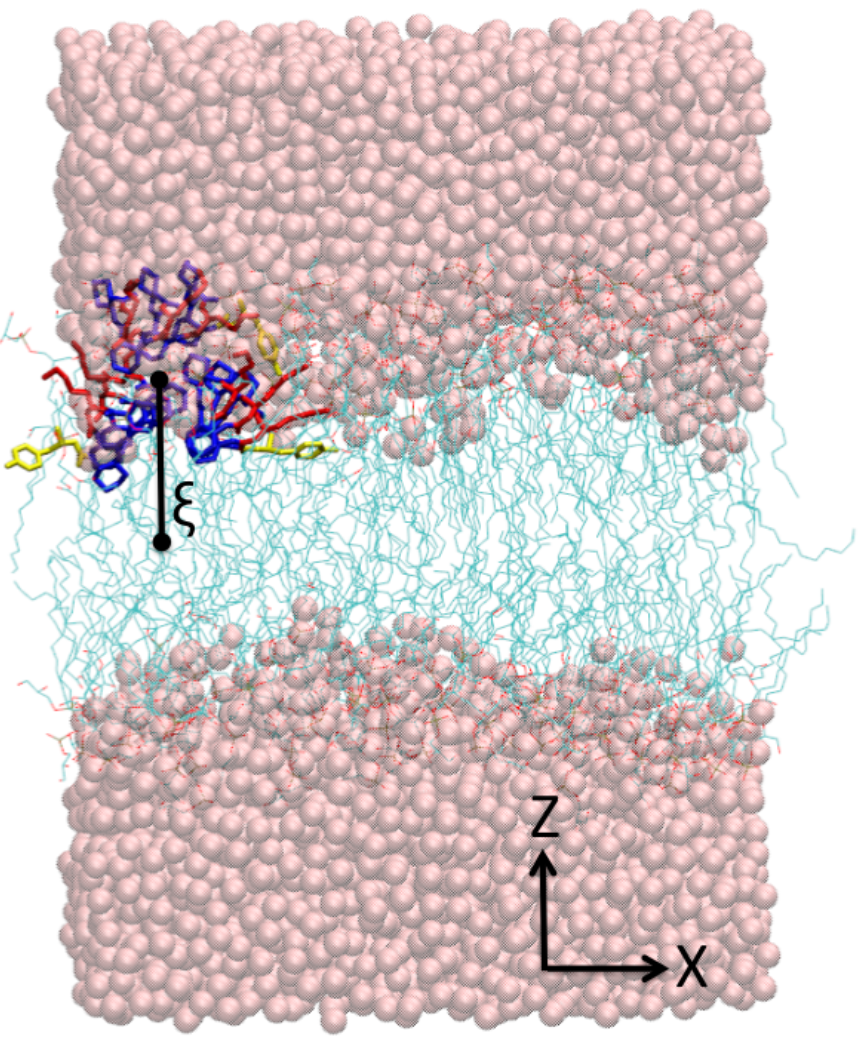
Schematic demonstration of the distance in Z direction between the center of mass co-ordinate of the peptide cluster and the membrane center as represented by the reaction coordinate *ξ*.

**Figure S2:**
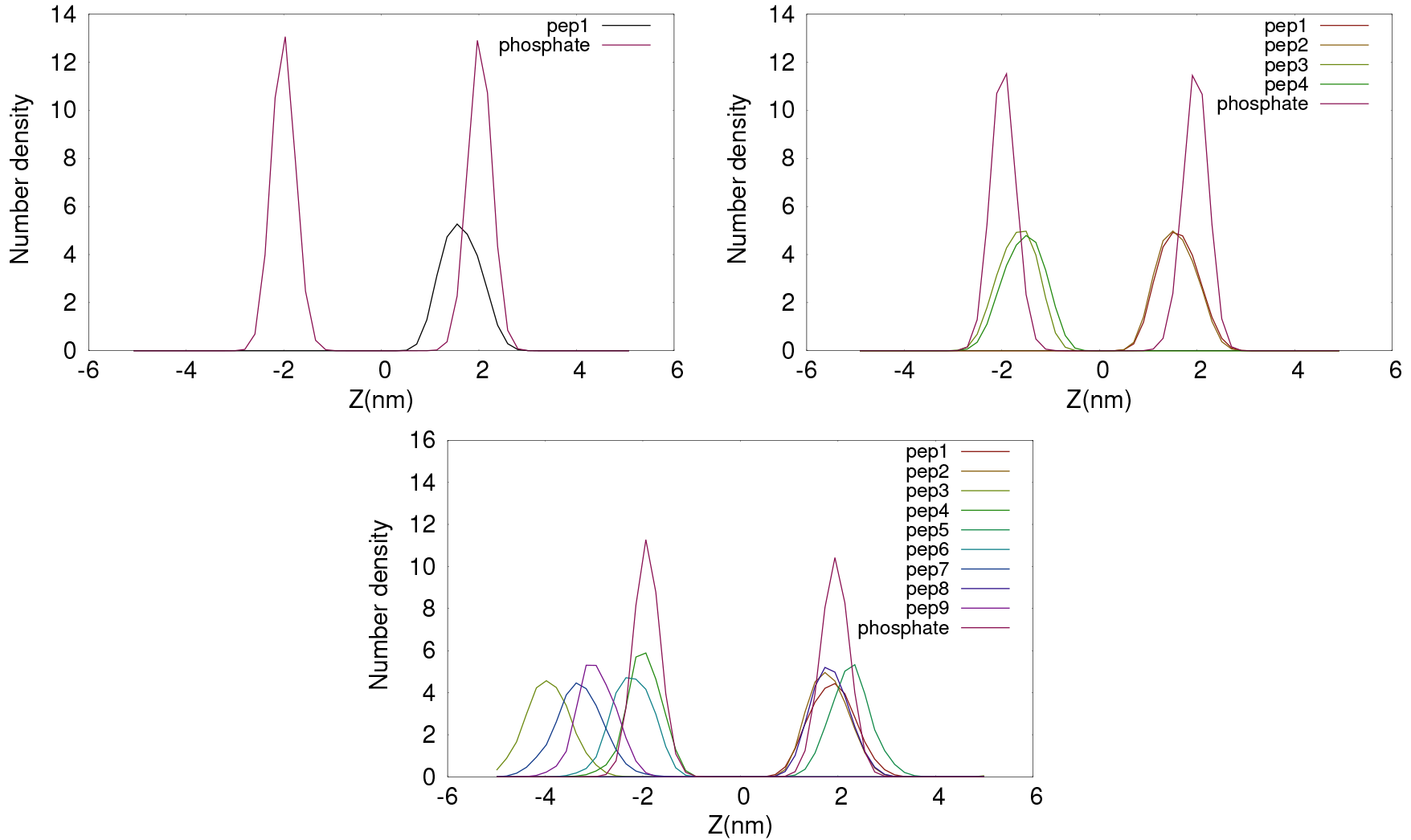
Density profile of Z direction of center of mass position of peptides relative to the membrane center of mass at low, medium and high P/L ratios in case of POPE/POPG lipid bilayer. Also plotted the density profile of phosphate groups of the lipid bilayers.

**Figure S3:**
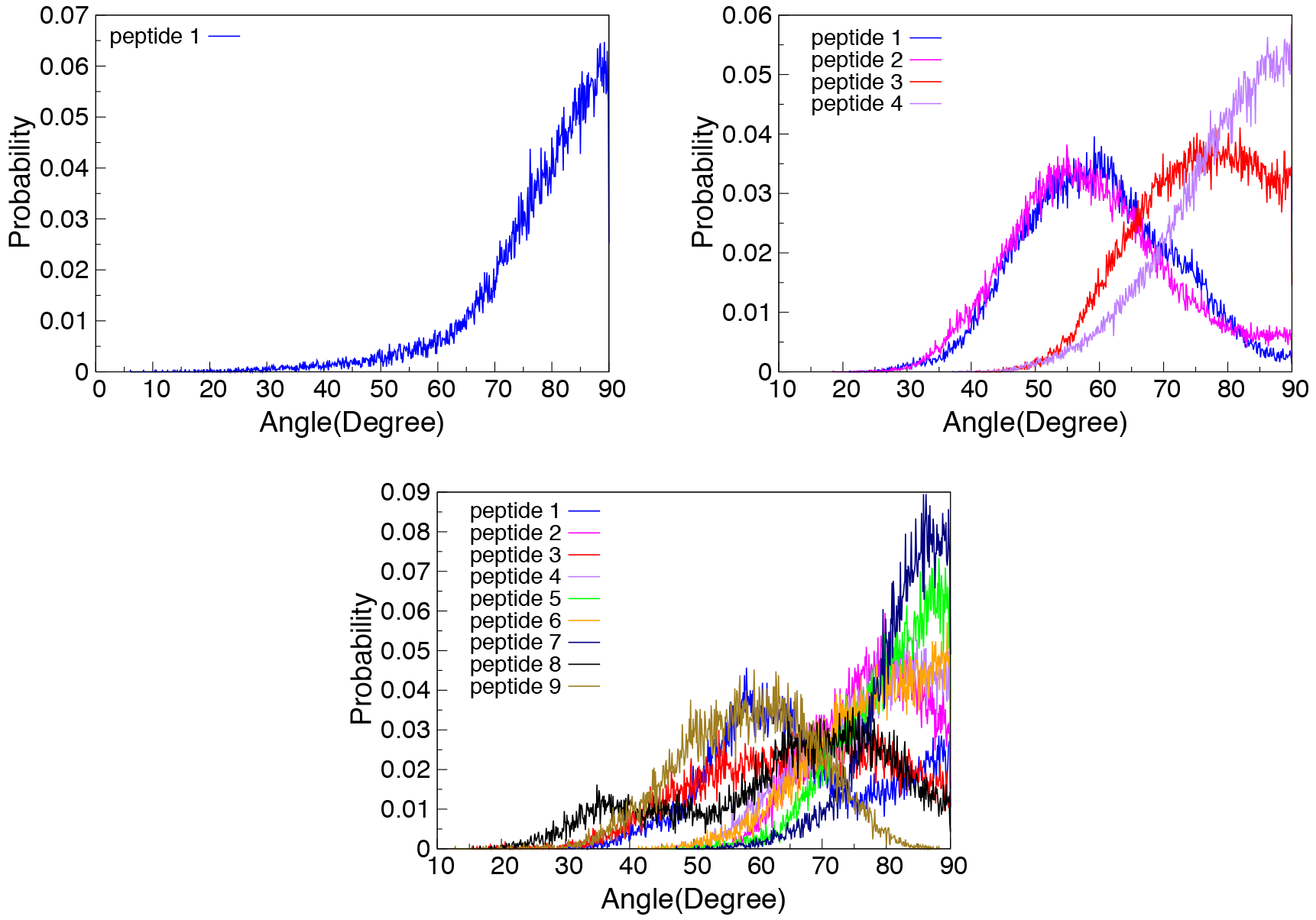
Probability distribution of angle of peptides with z-axis (normal to the membrane-surface) of DPPG bilayer at low, medium and high P/L ratios.

**Figure S4:**
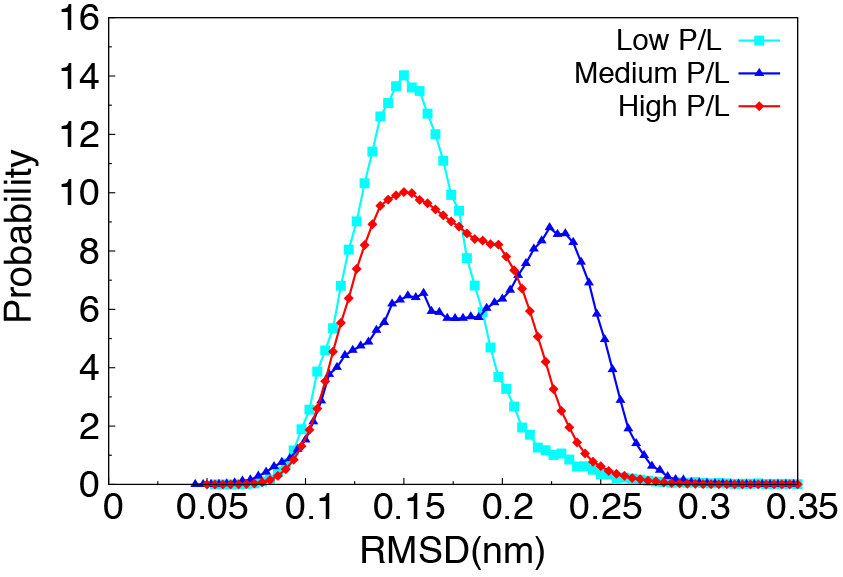
Probability distribution of root-mean-squared deviation of *β*-peptides from an ideal 14-helical structure at different P/L ratios.

**Figure S5:**
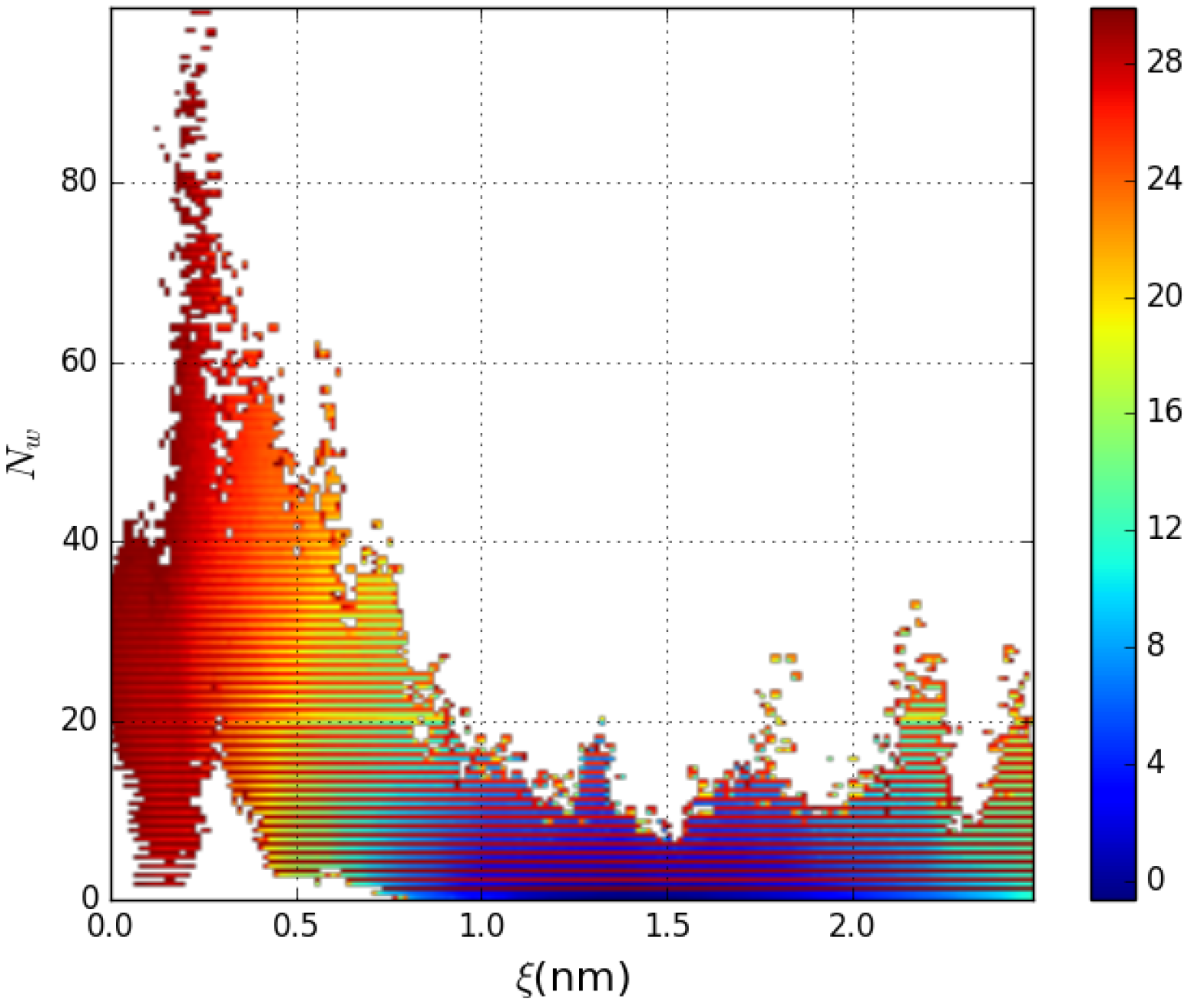
Two-dimensional projection of Free energy profile of the association of a three-peptide aggregate with the DPPG bilayer as a function of z-component distance of peptide-membrane center-of-mass separation (*ξ*) and number of water inside the membrane interior (*N*_*w*_). The free energy scale is reported in kcal/mol.

**Figure S6:**
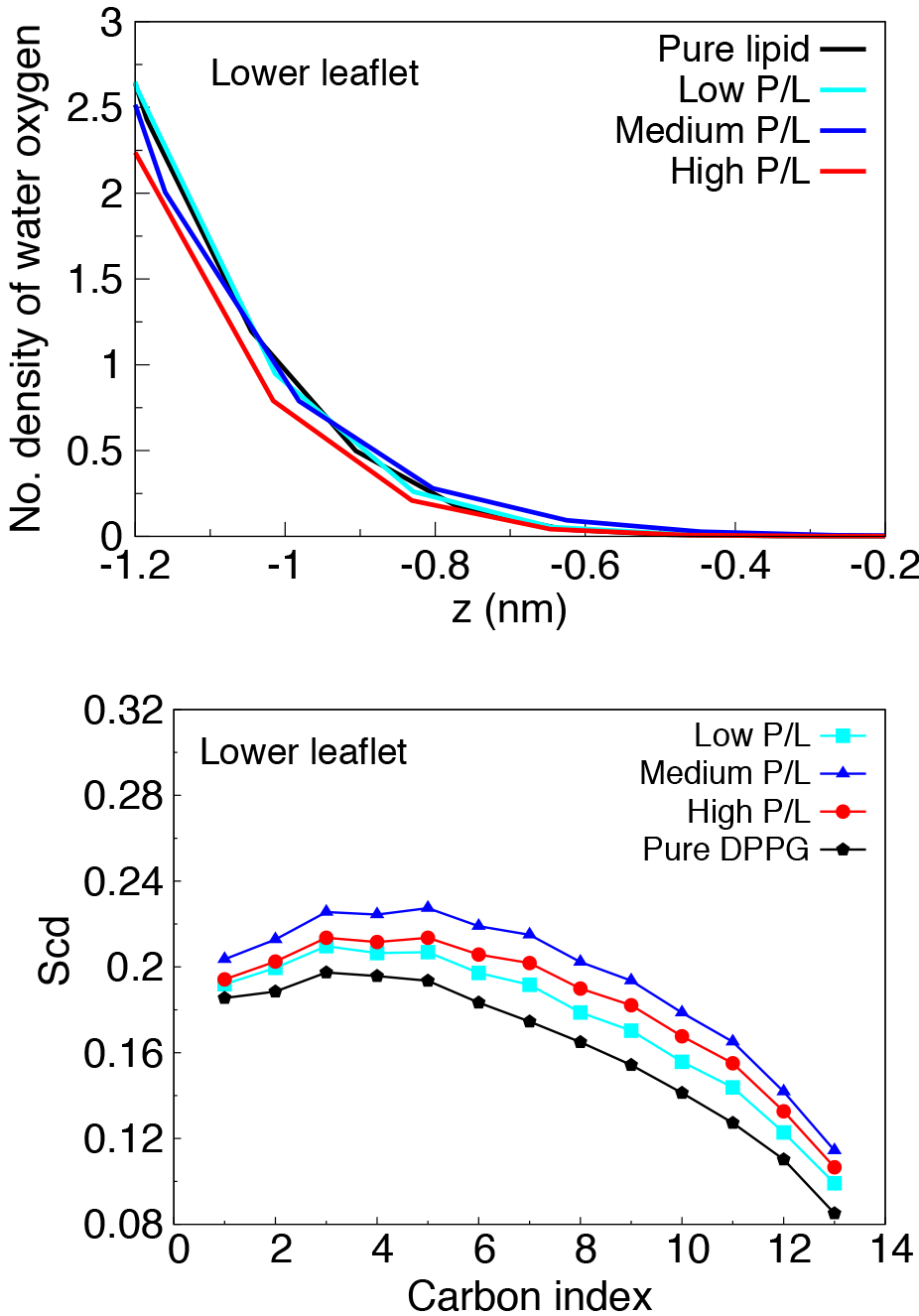
Bilayer property of Lower leaflet of the DPPG membranes : Top: The water-density profile of lower leaflets does not change appreciably with increase in P/L ratio. Bottom: The change in lipid order parameter values of lower leaflet of the bilayer with increasing P/L ratio is relatively smaller than that in upper leaflet.

**Figure S7:**
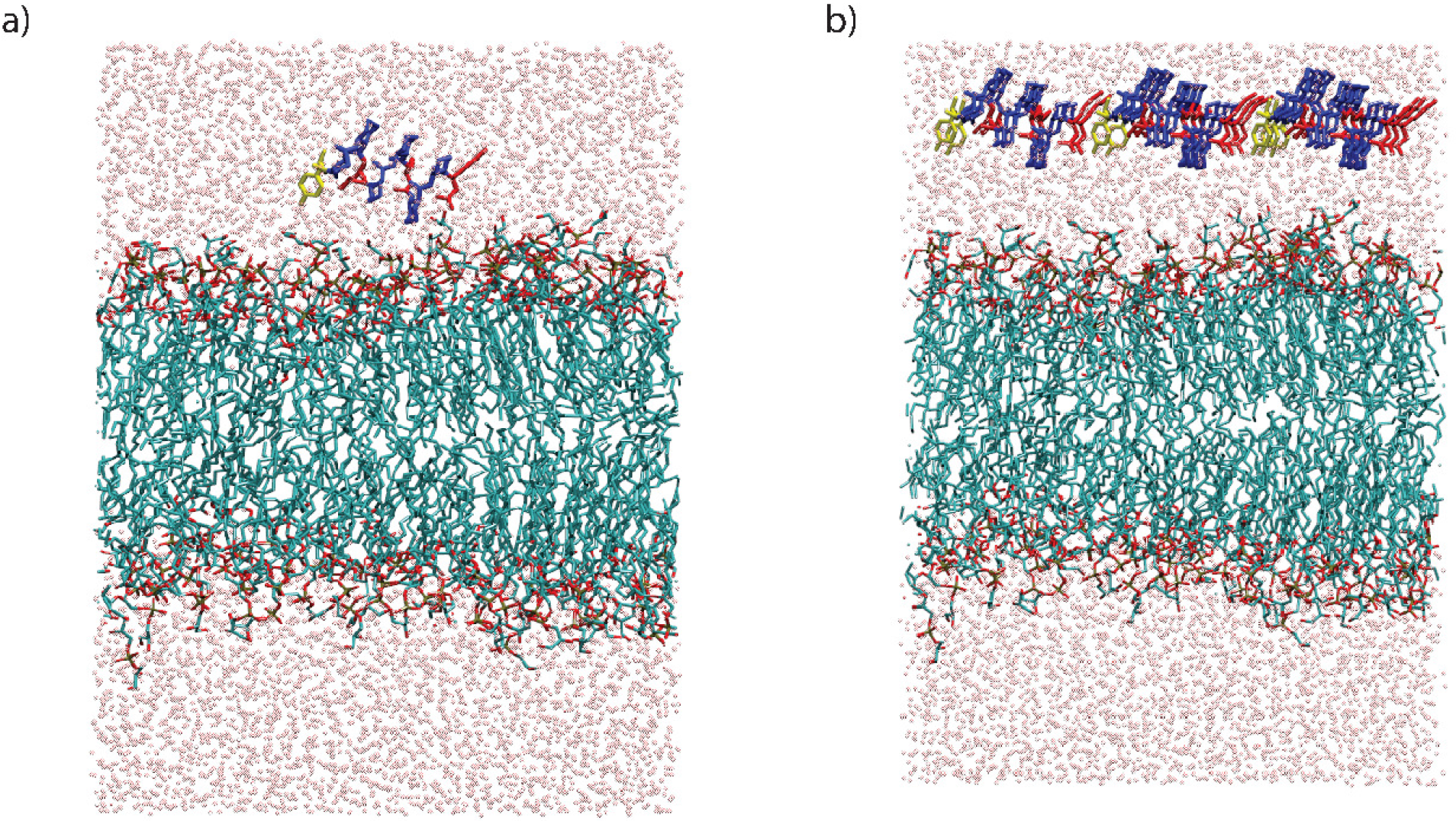
Snapshot of starting configurations for simulation of β-peptide(s) in presence of DPPG lipid bilayer: a) Initial configuration corresponding to simulation of single copy of β-peptide in the aqueous phase of the membrane, b) initial configuration corresponding to simulation of 9 copies β-peptides in aqueous phase of the DPPG membrane. All copies of β-peptides have been placed on the same side of the bilayer. Color code: lipid tails are cyan lines, head groups are red lines, water oxygen atoms are illustrated by pink spheres. The β-peptides are shown in licorice representation with hydrophobic ACHC residues in blue, β-lysine groups in red and β-tyrosines are in yellow.

## References

(1) Zassloff, M. Antimicrobial peptides of multicellular organisms. Nature 2002, 415, 389.

(2) Borgden, K. Antimicrobial peptides: pore formers or metabolic inhibitors in bacteria? Nature Rev. Microbiol. 2005, 3, 238.

(3) Yeaman, M.; Yount, N. Mechanisms of antimicrobial peptide action and resistance. Pharmacol.Rev. 2003, 55, 27.

(4) Shai, Y.; Oren, Z. From “carpet” mechanism to de-novo designed diastereomeric cell-selective antimicrobial peptides. Peptides 2001, 22, 1629.

(5) Porter, E. A.; Wang, X.; Lee, H.; Weisblum, B.; Gellman, S. H. Non-haemolytic β-amino-acid oligomers. Nature 2000, 404, 565–565.

(6) Liu, D.; DeGrado, W. F. De Novo Design, Synthesis, and Characterization of Antimicrobial β-Peptides. J. Am. Chem. Soc. 2001, 123, 7553–7559, PMID: 11480975.

(7) Chongsiriwatana, N. P.; Patch, J. A.; Czyzewski, A. M.; Dohm, M. T.; Ivankin, A.; Gidalevitz, D.; Zuckermann, R. N.; Barron, A. E. Peptoids that mimic the structure, function, and mechanism of helical antimicrobial peptides. Proc. Natl. Acad. Sci. 2008, 105, 2794–2799.

(8) Horne, W. S.; Boersma, M. D.; Windsor, M. A.; Gellman, S. H. Sequence-based design of alpha/beta-peptide foldamers that mimic BH3 domains. Angew.Chem. Int. Ed. 2008, 47, 2853–6.

(9) Frackenpohl, J.; Arvidsson, P. I.; Schreiber, J. V.; Seebach, D. The Outstanding Biological Stability of β- and γ-Peptides toward Proteolytic Enzymes: An In Vitro Investigation with Fifteen Peptidases. ChemBioChem 2001, 2, 445–455.

(10) Cheng, R. P.; Gellman, S. H.; Degrado, W. F. β-Peptides: From Structure to Function. Chem. Rev. 2001, 101, 3219.

(11) Gellman, S. H. Foldamers: A Manifesto. Acc. Chem. Res. 1998, 31, 173.

(12) Raguse, T. L.; Porter, E. A.; Gellman, S. H. Structure-Activity Studies of 14-Helical Antimicrobial β-Peptides:? Probing the Relationship between Conformational Stability and Antimicrobial Potency. J. Am. Chem. Soc. 2002, 124, 12774.

(13) Porter, E. A.; Weisblum, B.; Gellman, S. H. Mimicry of Host-Defense Peptides by Unnatural Oligomers:? Antimicrobial β-Peptides. J. Am. Chem. Soc. 2002, 124, 7324.

(14) Schmitt, M. A.; Weisblum, B.; Gellman, S. H. Interplay among Folding, Sequence, and Lipophilicity in the Antibacterial and Hemolytic Activities of α-β-Peptides. J. Am. Chem. Soc. 2007, 129, 417.

(15) Karlsson, A. J.; Pomerantz, W. L.; Gellman, S. H.; Palecek, S. P. Antifungal Activity from 14-Helical β-Peptides. J. Am. Chem. Soc. 2006, 128, 12630.

(16) Karlsson, A. J.; Pomerantz, W. L.; Neilsen,; J. K.; Gellman, S. H.; Palecek, S. P. Effect of Sequence and Structural Properties on 14-Helical 5-Peptide Activity against Candida albicans Planktonic Cells and Biofilms. ACS Chem. Biol. 2009, 4, 567.

(17) Shai, Y. Mode of action of membrane active antimicrobial peptides. Biopolymers 2002, 66, 236.

(18) Matsuzaki, K. Why and how are peptide?lipid interactions utilized for self-defense? Magainins and tachyplesins as archetypes. Biochim.Biophys.Acta 1999, 1462, 1.

(19) Leontiadou, H.; Mark, A. E.; Marrink, S. J. antimicrobial peptides in action. J. Am. Chem. Soc. 2006, 128, 12156.

(20) Sengupta, D.; Leontiadou, H.; Mark, A. E.; Marrink, S. J. Toroidal pores formed by antimicrobial peptides show significant disorder. Biochim.Biophys. Acta 2008, 1778, 2308.

(21) Perrin, B.; Pastor, R. Simulations of Membrane-Disrupting Peptides I: Alamethicin Pore Stability and Spontaneous Insertion. Biophys. J. 2016, 111, 1248–1257.

(22) Perrin, B.; Fu, R.; Cotten, M.; Pastor, R. Simulations of Membrane-Disrupting Peptides II: AMP Piscidin 1 Favors Surface Defects over Pores. Biophys. J. 2016, 111, 1258 – 1266.

(23) Zhu, X.; Konig, P.; Gellman, S. H.; Yethiraj, A.; Cui, Q. Establishing Effective Simulation Protocols for β and α-β-Peptides. II. Molecular Mechanical (MM) Model for a Cyclic β-Residue. J. Phys. Chem. B 2008, 112, 5439.

(24) Zhu, X.; Konig, P.; Hoffman, M.; Yethiraj, A.; Cui, Q. Establishing effective simulation protocols for β?and α-β?peptides. III. Molecular mechanical model for acyclic β?amino acids. J. Comput. Chem. 2010, 31, 2063.

(25) Pomerantz, W. C.; Abbott, N. L.; Gellman, S. H. Lyotropic Liquid Crystals from Designed Helical beta-Peptides. J. Am. Chem. Soc. 2006, 128, 8730–8731, PMID: 16819857.

(26) Pomerantz, W. C.; Yuwono, V. M.; Pizzey, C. L.; Hartgernik, J. D.; Abbott, N. L.; Gellman, S. H. Nanofibers and Lyotropic Liquid Crystals from a New Class of Self-Assembling β-Peptides. Angew.Chem. Int. Ed. 2008, 47, 1241.

(27) Miller, C. A.; Abbott, N. L.; Gellman, S. H.; de Pablo, J. J. Association of Helical β-Peptides and their Aggregation Behavior from the Potential of Mean Force in Explicit Solvent. Biophys. J. 2009, 96, 4349–4362.

(28) Dirac, A. D.; Moiset, G.; Mika, J. T.; Kocer, A.; Salvador, P.; Poolman, B.; Marrink, S. J.; Sengupta, D. The Molecular Basis for Antimicrobial Activity of Pore-Forming Cyclic Peptides. Biophys. J. 2011, 100, 2422.

(29) Morein, S.; Andersson, A.-S.; Rilfors, L.; Lindblom, G. Wild-type Escherichia coli Cells Regulate the Membrane Lipid Composition in a Window between Gel and Non-lamellar Structures. Journal of Biological Chemistry 1996, 271, 6801–6809.

(30) Pandit, K. R.; Klauda, J. B. Membrane models of E. coli containing cyclic moieties in the aliphatic lipid chain. Biochim,. Biophys. Acta - Biomembranes 2012, 1818, 1205 – 1210.

(31) Torrent, M.; Pulido, D.; Nogues, M. V.; Boix, E. Exploring New Biological Functions of Amyloids: Bacteria Cell Agglutination Mediated by Host Protein Aggregation. PLOS Pathogens 2012, 8, 1–8.

(32) Kumar, D. K. V.; Choi, S. H.; Washicosky, K. J.; Eimer, W. A.; Tucker, S.; Ghofrani, J.; Lefkowitz, A.; McColl, G.; Goldstein, L. E.; Tanzi, R. E.; Moir, R. D. Amyloid-βpeptide protects against microbial infection in mouse and worm models of Alzheimer’s disease. Science Translational Medicine 2016, 8, 340ra72–340ra72.

(33) Miller, C. A.; Abbott, N. L.; Gellman, S. H.; de Pablo, J. J. Mechanical Stability of Helical β-Peptides and a Comparison of Explicit and Implicit Solvent Models. Biophys. J. 2008, 95, 3123.

(34) Irudayam, S. J.; Berkowitz, M. L. Influence of the arrangement and secondary structure of melittin peptides on the formation and stability of toroidal pores. Biochim. Biophys. Acta - Biomembranes 2011, 1808, 2258 – 2266.

(35) Irudayam, S. J.; Pobandt, T.; Berkowitz, M. L. Free Energy Barrier for Melittin Reorientation from a Membrane-Bound State to a Transmembrane State. J. Phys. Chem. B 2013, 117, 13457–13463, PMID: 24117276.

(36) Baul, U.; Kuroda, K.; Vemparala, S. Interaction of multiple biomimetic antimicrobial polymers with model bacterial membranes. J. Chem. Phys. 2014, 141, 084902.

(37) Yang, L.; Gordon, V.; Mishra, A.; Som, A.; Purdy, K.; Davis, M.; Tew, M.; Wong, G. Synthetic Antimicrobial Oligomers Induce a Composition-Dependent Topological Transition in Membranes. J.Am.Chem.Soc. 2007, 129, 12141.

(38) Lee, M. W.; Chakraborty, S.; Schmidt, N. W.; Murgai, R.; Gellman, S. H.; Wong, G. C. Two interdependent mechanisms of antimicrobial activity allow for efficient killing in nylon-3-based polymeric mimics of innate immunity peptides. Biochim. Biophys. Acta-Biomembranes 2014, 1838, 2269 – 2279, Interfacially active peptides and proteins.

(39) Choi, H.; Chakraborty, S.; Liu, R.; Gellman, S. H.; Weisshaar, J. C. Single-Cell, Time-Resolved Antimicrobial Effects of a Highly Cationic, Random Nylon-3 Copolymer on Live Escherichia coli. ACS Chem. Biol. 2016, 11, 113–120, PMID: 26493221.

(40) Liu, R.; Chen, X.; Falk, S. P.; Masters, K. S.; Weisblum, B.; Gellman, S. H. Nylon-3 Polymers Active against Drug-Resistant Candida albicans Biofilms. J. Am. Chem. Soc. 2015, 137, 2183–2186, PMID: 25650957.

(41) Chakraborty, S.; Liu, R.; Hayouka, Z.; Chen, X.; Ehrhardt, J.; Lu, Q.; Burke, E.; Yang, Y.; Weisblum, B.; Wong, G. C. L.; Masters, K. S.; Gellman, S. H. Ternary Nylon-3 Copolymers as Host-Defense Peptide Mimics: Beyond Hydrophobic and Cationic Subunits. J. Am. Chem. Soc. 2014, 136, 14530–14535, PMID: 25269798.

(42) Last, N. B.; Schlamadinger, D. E.; Miranker, A. D. A common landscape for membrane-active peptides. Protein Science 2013, 22, 870–882.

(43) Dufourcq, J.; Faucon, J.-F.; Fourche, G.; Dasseux, J.-L.; Maire, M. L.; Gulik-Krzywicki, T. Morphological changes of phosphatidylcholine bilayers induced by melittin: vesicularization, fusion, discoidal particles. Biochim. Biophys. Acta - Biom,em,-branes 1986, 859, 33 – 48.

(44) Therrien, A.; Lafleur, M. Melittin-Induced Lipid Extraction Modulated by the Methylation Level of Phosphatidylcholine Headgroups. Biophys. J. 2015, 110, 400–410.

(45) Therrien, A.; Fournier, A.; Lafleur, M. Role of the Cationic C-Terminal Segment of Melittin on Membrane Fragmentation. J. Phys. Chem. B 2016, 120, 3993–4002.

(46) Jo, S.; Kim, T.; Iyer, V. G.; Im, W. CHARMM-GUI: A web-based graphical user interface for CHARMM. J. Comput. Chem. 2008, 29, 1859–1865.

(47) Best, R. B.; Zhu, X.; Shim, J.; Lopes, P. E. M.; Mittal, J.; Feig, M.; MacKerell, A. D. Optimization of the Additive CHARMM All-Atom Protein Force Field Targeting Improved Sampling of the Backbone œï, œà and Side-Chain œá1 and œá2 Dihedral Angles. J.Chem.Theo. Comput. 2012, 8, 3257–3273, PMID: 23341755.

(48) Mondal, J.; Zhu, X.; Cui, Q.; Yethiraj, A. Sequence-dependent interaction of β-peptides with membranes. J.Phys.Chem.B 2010, 114, 13585.

(49) MacKerell, A. D. et al. All-Atom Empirical Potential for Molecular Modeling and Dynamics Studies of Proteins. J. Phys. Chem. B 1998, 102, 3586–3616, PMID: 24889800.

(50) Tieleman, D. P.; Marrink, S.-J. Lipids Out of Equilibrium: Energetics of Desorption and Pore Mediated Flip-Flop. J. Am. Chem. Soc. 2006, 128, 12462–12467, PMID: 16984196.

(51) Bennett, W. F.; Sapay, N.; Tieleman, D. P. Atomistic Simulations of Pore Formation and Closure in Lipid Bilayers. Biophys. J. 2014, 106, 210 – 219.

(52) DomanÑski, J.; Hedger, G.; Best, R. B.; Stansfeld, P. J.; Sansom, M. S. P. Convergence and Sampling in Determining Free Energy Landscapes for Membrane Protein Association. J. Phys. Chem. B 2017, 121, 3364–3375, PMID: 27807980.

(53) A. Grossfield, WHAM: the weighted histogram analysis method, version 2.0. http://membrane.urmc.rochester.edu/content/wham.

(54) Awasthi, N.; Hub, J. S. Simulations of Pore Formation in Lipid Membranes: Reaction Coordinates, Convergence, Hysteresis, and Finite-Size Effects. J. Chem.Theory. Comput. 2016, 12, 3261–3269, PMID: 27254744.

(55) Hess, B.; Kutzner, C.; Van Der Spoel, D.; Lindahl, E. GROMACS 4: algorithms for highly efficient, load-balanced, and scalable molecular simulation. J.Chem.Theo. Comput. 2008, 4, 435–447.

(56) Nosé, S. A molecular dynamics method for simulations in the canonical ensemble. Mol. Phys. 1984, 52, 255.

(57) Hoover, W. Canonical dynamics: equilibrium phase-space distributions. Phys. Rev. A 1985, 31, 1695.

(58) Parrinello, M.; Rahman, A. Polymorphic transitions in single crystals: A new molecular dynamics method. J. App. Phys. 1981, 52, 7182–7190.

(59) Pail, S.; Hess, B. A flexible algorithm for calculating pair interactions on {SIMD} architectures. Comput. Phys. Comm.s 2013, 184, 2641 – 2650.

(60) Hess, B.; Bekker, H.; Berendsen, H. J. C.; Fraaije, J. G. E. M. LINCS: A linear constraint solver for molecular simulations. J.Comput.Chem. 1997, 18, 1463–1472.

(61) Miyamoto, S.; Kollman, P. Settle: An analytical version of the SHAKE and RATTLE algorithm for rigid water models. J. Comput. Chem. 1992, 13, 952–962.

(62) Marrink, S.-J.; de Vries, A.; Mark, A. Coarse Grained Model for Semiquantitative Lipid Simulations. J.Phys.Chem.B 2004, 108, 750.

(63) Allen, W. J.; Lemkul, J. A.; Bevan, D. R. GridMAT-MD: A grid-based membrane analysis tool for use with molecular dynamics. J. Comput. Chem. 2009, 30, 1952–1958.

